# mRNA psi profiling using nanopore DRS reveals cell type-specific pseudouridylation and translational regulation

**DOI:** 10.1101/2024.05.08.593203

**Authors:** Caroline A. McCormick, Michele Meseonznik, Yuchen Qiu, Oleksandra Fanari, Mitchell Thomas, Yifang Liu, Dylan Bloch, Isabel N. Klink, Miten Jain, Meni Wanunu, Sara H. Rouhanifard

## Abstract

Pseudouridine (psi) is one of the most abundant human mRNA modifications yet it’s functional impact on translation has remained unclear. Using direct RNA nanopore sequencing coupled with our Mod-*p* ID analytical framework, we mapped psi at single-nucleotide resolution across six immortalized human cell lines derived from diverse tissue types. Psi sites identified by nanopore sequencing were cross-validated using Illumina-based methods, confirming both positional accuracy and reproducibility. Unlike prior short-read approaches, nanopore sequencing provided the unique ability to quantify relative occupancy at each site and to detect multiple modifications on the same RNA molecule, revealing combinatorial modification patterns that cannot be captured otherwise. Integrating these psi maps with matched proteomic and ribosome profiling datasets, we find that psi modulates translation through two mechanistic modes: (i) single high-occupancy psi sites enhance translational efficiency and protein output, whereas (ii) clustered psi modifications promote ribosome pausing, decoupling translation efficiency from protein yield. This integrative, multi-omics framework provides a quantitative model of how psi stoichiometry and distribution along transcripts shape ribosome dynamics and proteome composition across human cell types.

## Introduction

Pseudouridine (psi) is an RNA modification resulting from the isomerization of uridine by the psi synthase family of enzymes (PUS). Psi is found in multiple classes of RNAs, including noncoding RNAs (ncRNAs), and messenger RNA (mRNAs). On mRNAs, psi has been implicated in modulating translational fidelity(1, 2), structural stability(3), and pre-mRNA processing(4). Remarkably, studies demonstrate that psi within coding sequences can promote ribosomal readthrough of premature termination codons (PTCs), effectively restoring full-length protein synthesis in transcripts that would otherwise produce truncated, non-functional proteins(5). Notably, this effect can be recapitulated through programmable pseudouridylation, in which psi is intentionally installed at specific codons to suppress nonsense mutations— highlighting a causable and tunable link between psi placement and translational outcomes(6). While these findings highlight one mechanism by which psi can influence decoding, they also underscore a broader question: how else does psi shape translation across the transcriptome? Specifically, the extent to which psi stoichiometry, positional context and clustering alter ribosome dynamics and protein synthesis remains largely unexplored.

Recent work has revealed that psi is not uniformly distributed but instead exhibits marked cell type-specific and state-dependent variation. In neuronal models, we found that psi landscapes are partially remodeled during differentiation, with certain sites remaining highly static across cell states while others are highly plastic, responding to changes in differentiation status or environmental perturbation(7). In particular, comparisons of undifferentiated, differentiated and lead-treated SH-SY5Y cells revealed subsets of psi sites that were selectively induced or lost under stress conditions, suggesting that psi may contribute to both developmental and adaptive translational programs(7). In immune cells, psi occupancy differs between immortalized and primary contexts, consistent with transformation-dependent reprogramming of RNA modification profiles. Together these findings suggest that psi is a regulated and context specific mark with the potential to shape translation in ways that depend on cell identity, environmental cues and physiological state.

To discern the role of psi in post-transcriptional regulation, we need a comprehensive map that outlines psi positions and occupancy levels across diverse human cell types. Previous approaches have identified psi positions (8–10) and the relative psi-to-uridine fractions in mRNA (8, 9) across cell lines. However, these methods rely on either *N*-cyclohexyl-*N*′-(2-morpholinoethyl)carbodiimide methyl-*p*-toluenesulfonate (CMC) to block the progress of reverse transcriptase during library preparation(11, 12) or bisulfite detection which results in a systematic deletion signature during reverse transcription(8, 10, 13). While powerful, these cDNA-based techniques are limited by amplification bias (14) and can yield false positives when the detected psi has neighboring uracil sites(8). Importantly, because these methods rely on short-read sequencing, they cannot determine whether multiple modifications occur on the same RNA molecule, precluding the analysis of combinatorial or co-occurring modification patterns that may influence translation.

Nanopore Direct RNA Sequencing (DRS) is currently the only method to detect native human RNA without conversion to cDNA. Nanopore DRS employs a motor protein sequencing adapter that allows for the voltage-driven translocation of RNA through a biological nanopore. A characteristic disruption of the ionic current is associated with each canonical nucleotide during basecalling allowing for the conversion of a strand’s current signal to a corresponding local sequence. Because an RNA strand is passed through the pore (i.e., not a converted cDNA strand), chemical modifications to the RNA nucleobases often produce aberrations in the expected ionic current levels(15). DRS has been used to map several mRNA modifications to date, including psi(16–18), *N*^6^-methyladenosine (m^6^A)(19–24), and inosine (I)(25) across the human transcriptome.

Psi sites can be detected from DRS as sequence -specific U-to-C basecalling errors. Using our Mod-p ID framework(16, 26). These errors are compared against an unmodified in vitro transcribed (IVT) control(27) to confidently assign psi sites transcriptome-wide. Importantly, psi calls widentified by Mod-p ID were cross validated across platforms. Unlike short-read approaches, nanopore DRS enables quantification of relative occupancy at each site and identification of multiple modifications on the same RNA molecule, revealing patterns of psi spatial organization—from isolated high-occupancy sites to densely clustered regions—on individual transcripts(28). These features allow DRS to capture both the stoichiometry and positional context of psi modifications that together shape translational outcomes.

Here, we apply Mod-*p* ID to perform a comparative, transcriptome-wide analysis of psi in six diverse, immortalized human cell lines: A549 (lung), HeLa (cervix), HepG2 (liver), Jurkat (T cells), NTERA (testes), SH-SY5Y (neuron-like). We quantify relative psi occupancy across cell types, examine its relationship to psi synthase expression, and assess how psi correlates with protein abundance using quantitative proteomics. Building on these data, we integrate proteomics and ribosome profiling (Ribo-seq) to measure translational efficiency (TE) and ribosome pausing, providing a mechanistic model of how psi regulates translation through two distinct modes: single high-occupancy sites that enhance translation and clustered psi sites that promote ribosome pausing. Together, this integrated multi-omic framework reveals how psi stoichiometry and positional organization coordinate translational regulation in human cells and may inform future strategies for programmable therapeutic pseudouridylation.

## Materials & Methods

### Accessing publicly available data sets

Nanopore DRS raw fast5s for all IVTs (RNA002) were sourced from NIH NCBI SRA BioProject accession PRJNA947135(27). Nanopore DRS raw fast5 files for native HeLa replicates (RNA002) were sourced from NIH NCBI SRA BioProject accession PRJNA777450(16). Nanopore DRS raw fast5s for native SH-SY5Y replicates (RNA002) were sourced from NIH NCBI SRA BioProject accession PRJNA1092333(7). Nanopore DRS fastq files for A549 cells and corresponding IVT control (RNA004) can be found in NIH NCBI SRA BioProject accession PRJNA1329214.

### Cell culture

HeLa, HepG2, A549, and NTERA-2 cells were cultured in DMEM (Gibco, 10566024); Jurkat cells were cultured with RPMI 1640 (Gibco™ 11875093); Jurkat cells were cultured in RPMI 1640 (Cat No. 11-875-093) and SH-SY5Y cells were cultured in 1:1 EMEM: F12 (Fisher Scientific, 50-983-283, Cytiva, SH3002601). All media was supplemented with 10% Fetal Bovine Serum (Fisher Scientific, FB12999102) and 1% Penicillin-Streptomycin (Lonza,17602E). Cells were cultured at 37°C with 5% CO_2_ in 10 cm tissue culture dishes and allowed to reach ∼80% confluence.

### Total RNA extraction and poly(A) selection

Total RNA extraction from cells and Poly(A) selection was performed using a previously established protocol(16). Six confluent 10 cm cell culture dishes were washed with ice-cold PBS, lysed with TRIzol (Invitrogen,15596026) at room temperature, and transferred to an RNAse-free microcentrifuge tube. Chloroform was added to separate the total RNA in the aqueous supernatant from the organic phase following centrifugation. The aqueous supernatant was extracted and transferred to a new RNAse-free microcentrifuge tube. An equal volume of 70% absolute ethanol was added. PureLink RNA Mini Kit (Invitrogen, 12183025) was used to purify the extracted total RNA following the Manufacturer’s protocol. Total RNA concentration was measured using the Qubit™ RNA High Sensitivity (HS) assay (Thermo Fisher, Q32852). Poly(A) selection was performed using NEBNext Poly(A) mRNA Magnetic Isolation Module (NEB, E7490L) according to the Manufacturer’s protocol. The isolated poly(A) selected RNA was eluted from the beads using Tris buffer. The poly(A) selected RNA concentration was measured using the same Qubit™ assay listed above.

### *In vitro* transcription and polyadenylation

The protocol for IVT, capping, and polyadenylation is described previously(16).

### Sequencing, basecalling, and alignment procedure

Cell line replicates were prepared for Nanopore Direct RNA sequencing (DRS) following the ONT SQK-RNA002 kit protocol, including the reverse transcription step. RNA sequencing on the MinION and PrometheION platforms was performed using ONT R9.4.1 (FLO-MIN106) and (PRO-002). Jurkat and NTERA cells were sequenced on the PromethION while all other cell lines were sequenced on the MinION.

DRS runs were basecalled with Guppy v6.4.2 using the high accuracy model and the default basecalling quality score filter of Q ≥7(35). Basecalled read replicates for a cell line were merged and then aligned with minimap2 to the GRCh38.p10 reference genome for downstream analysis.

### Genomic DNA extraction and Sanger sequencing

To analyze putative SNVs, we performed Sanger sequencing on genomic DNA (gDNA). gDNA extraction was performed using a Monarch Genomic DNA Purification Kit (NEB, T3010S) following the Manufacturer’s protocol. PCR primers were designed to amplify ∼300 nt regions surrounding the detected Ψ positions using Primer-BLAST with default settings (**Supplementary Table S1**). Using the Manufacturer’s protocol, the PCR reaction was set up with Q5 polymerase (NEB, M0491L). Thermocycling conditions were as follows: initial denaturation at 98°C for 30 seconds, 25 cycles of 98°C for 10 seconds, then 63°C for 20 seconds and 72°C for 15 seconds, final extension at 72°C for 2 minutes, and holding at 10°C. PCR products were purified using a Monarch PCR & DNA Cleanup Kit (NEB, T1030S) following the Manufacturer’s protocol. The concentration of eluted DNA was determined using a Nanodrop spectrophotometer. The purified PCR products were imaged on a 2% agarose TBE gel to confirm specific amplification. Samples were sent to Quintara Biosciences for SimpliSeq™ Sanger sequencing. Results are shown in **Supplementary Figure S1**.

### Immunofluorescence staining and analysis of PUS7 and TRUB1

All six cell lines were seeded in Lab Tek 8-well chambers (Thermo Scientific™ Cat No.155409PK). A549 and HeLa cells were seeded at 30k cells/well, HepG2 and NTERA cells were seeded at 60k cells/well, and SHSY5Y cells were seeded at 100k cells/well. A549, Hela, HepG2, and NTERA cell lines were cultured in 300uL DMEM (Gibco Cat No. 10566-016) and SHSY5Y cells were cultured in 300uL 1:1 EMEM:F12 media (Quality Biological Cat No. 112-018-131, Gibco Cat No. 11765054). After 24 hours, half the media in each well was removed and replenished with equal volume 4% formaldehyde. Following 2-minute incubation at room temperature, the 2% formaldehyde was aspirated and replaced with 300 µL of 4% formaldehyde. Cells incubated at room temperature for 10 minutes before they were washed twice for 5 minutes and stored in PBS at 4°C.

Cells were removed from 4°C and all PBS was aspirated. Each well was permeabilized with 300 µL of 0.1% PBS-Triton X-100 for 10 minutes at room temperature. The entire solution was aspirated and each well was blocked with 300uL of 2% BSA in 0.1% PBS-Triton X-100. Cells were incubated at room temperature for one hour and washed 3 times with 0.1% PBS-Tween20. Two wells in each plate were treated with 300 µL of 400 µg/µL anti-PUS7 antibodies (Prestige Antibodies Cat No. HPA024116) or 200ug/uL anti-TRUB1 antibodies (Protein Tech Cat No. 12520-1-AP) in 1% BSA in PBS-Triton. Each well treated with a primary antibody was also stained with 5 µg/mL GAPDH conjugated antibodies (Invitrogen Catalog No. MA5-15738-D488). 2 wells in each plate were left as no primary antibody controls. Cells were incubated overnight in the dark at 4°C and washed once with 0.1% PBS-Tween20. Secondary antibody staining was performed with 1:500 Alexa Fluor® 594 AffiniPure™ Alpaca Anti-Rabbit IgG (H+L) antibodies (Cat No. 611-585-215). Following 1 hour incubation at room temperature the cells were washed once and stained with 167 ng/mL DAPI for 20 minutes. Cells were then washed and stored in 2X SSC.

The analysis of cell staining was conducted using CellProfiler(29). The details are in the CellProfiler project file (IF_analysis.cpproj). Briefly, the DAPI, GFP and Alexa594 channels of each image containing many cells were used for analysis. The DAPI channel was used to segment the nuclei by the typical diameter of the object. The GFP channel was used to segment the cells by adaptive minimum cross-entropy thresholding on the logarithm of intensity. The fluorescence intensities of the GFP and Alexa594 channels were measured based on the mean intensity of each channel in the cells. For the IF analysis, the mean intensity of the Alexa594 channel is divided by the mean intensity of the GFP channel to obtain the fold change for both TRUB1 and PUS7 targets. Ten fluorescence microscopy images from two biological replicates were obtained for the analysis.

### Protein Extraction and LC-MS/MS

Cell lysis buffer was prepared using 8M urea (Fisher, Cat no: U15-500), 40mM Tris-HCl, pH 7.0 (Fisher, Cat no: BP1756-100), and Pierce Protease Inhibitor Tablets (Thermo Scientific, Cat no: PIA32963). Cells from six cell lines—A549 (3.492 × 10^7 cells), HeLa (3.240 × 10^7 cells), HepG2 (2.946 × 10^7 cells), Jurkat (7.070 × 10^7 cells), SH-SY5Y (7.236 × 10^7 cells), and NTERA-2 (1.392 × 10^7 cells)—were lysed using this buffer. The lysates were incubated on ice for 30 minutes and then centrifuged at 20,000 x g for 15 minutes at 4°C. Supernatants were collected, and protein concentrations were measured using the Qubit Protein Assay Kit (Invitrogen by Thermo Fisher Scientific, Cat no: Q33211).

100 µg of protein per sample from three biological replicates per group was taken for proteomics analysis. Samples were reduced with 10 mM TCEP at 56°C for 60 mins and alkylated using 20 mM iodoacetamide at RT for 30 mins. Before digestion, proteins were precipitated with acetone. The protein pellet was re-suspended in 100 mM Triethylamonium bicarbonate (TEAB) and digested with MS-grade trypsin (Pierce TM) overnight at 37°C. Peptide digest was quantified using Thermo Scientific™ Pierce™ Quantitative Colorimetric Peptide Assay (Product No. 23275). An equal amount of peptide was labeled with isobaric TMT reagents Thermo Scientific™ TMT10plex (Pierce; Rockford, IL, USA) according to manufacturer protocols (Thermo Scientific TM). A small fraction of the sample from each isobaric tag channel was pooled and analyzed on LC-MS to obtain the median for all the reporter ion intensities. TMT-labeled peptides were pooled and cleaned with Pierce™ C18 spin columns (Thermo Scientific™) and were re-suspended in 2% acetonitrile (ACN) and 0.1% formic acid (FA). 1 μg of multiplexed sample was loaded onto in-house pull tip 75µm × 20 cm C18 ReproSil-Pur 120 1.9 µm LC Column, then separated with a Thermo RSLC Ultimate 3000 (Thermo Scientific™) with a 160 min gradient of 2–35% solvent B (0.1% FA in ACN) at 200 nL/min and 25oC with a 220 min total run time. Eluted peptides were analyzed by a Thermo Orbitrap Q Exactive (Thermo Scientific™) mass spectrometer in a data-dependent acquisition mode. A full survey scan MS (from m/z 350–1500) was acquired in the Orbitrap with a resolution of 70,000. The AGC target for MS1 was set as 3 × 106, and the ion filling time was set as 50 ms. The most intense 15 ions with charge states 2-6 were isolated and fragmented using HCD fragmentation with 33 % normalized collision energy and detected at a mass resolution of 35,000 at 200 m/z. The AGC target for MS/MS was set as 1 × 105, and the ion filling time was set as 120 ms. Dynamic exclusion was set for 30 s with a 0.7 m/z isolation window.

### Proteomics Analysis

Raw LC-MS/MS data were loaded into MaxQuant(30) v2.4.2.0 and the peptides were identified with the built-in Andromeda search engine using a FASTA file containing all entries from the SwissProt Homo sapiens (human) database (20,218 proteins) supplemented with common contaminants.

For all searches, carbamidomethylated cysteine was set as a fixed modification and oxidation of methionine and N-terminal protein acetylation as variable modifications. Trypsin/P was specified as the proteolytic enzyme. Precursor tolerance was set to ±10 ppm, and fragment ion tolerance to ±20ppm. For modified peptides, we used the default cutoffs of at least 40 for the Andromeda score and 6 for the delta score. The false discovery rate was set at <1% for peptide spectrum matches and protein group identification employing a target-decoy approach.

Reporter ion isotropic distributions from the TMT product data sheet were input into the software to correct for labeling. The peptides were filtered to have <0.05 PEP (posterior error probability) and all the contaminants were excluded. We filtered the peptides to be present (Reporter ion intensity > 0) in all replicates of each cell line. The median reporter ion intensities were then calculated to account for the relative lengths of the proteins for quantification. The MaxQuant output can be found in **Supplementary Table S2**. Normalized protein abundance and corresponding mRNA TPMs can be found in **Supplementary Table S3**.

### Bootstrapping

Since we compared MinION and PromethION runs, which generate substantially different numbers of reads, we performed computational resampling to generate ten *in silico* replicates for the native mRNA with 1.2M direct reads, each randomly sampled from the base called FASTQ files for each cell line. To account for the reduction in the total number of reads, we relaxed the threshold from 30 direct reads to 10 direct reads at a query position.

### Effect size analysis of pseudouridine site number on expression and Translation Efficiency Analysis

To assess the impact of conserved ψ site number on gene expression, transcripts were divided into three groups based on the number of psi-sites annotated in Supplementary Table S7: 1) transcripts with no psi sites, 2) transcripts with one ψ site, and 3) transcripts with more than two ψ sites.

Protein abundance values were obtained from the median protein abundance estimates in Supplementary Table S3. mRNA expression levels were measured as transcripts per million (TPM) from the same table. Translation efficiency (TE) values were calculated from publicly available ribosome profiling (Ribo-seq) and RNA-seq datasets obtained from GEO (HeLa: GSE21992, GSE79664, GSE143301, GSE188692, SRA099816; A549: GSE82232, GSE101760; HepG2: GSE125757, GSE174419; SH-SY5Y: GSE148827, GSE155727). For each transcript, TE was defined as the ratio of ribosome-protected fragment counts to RNA-seq counts, as compiled in Supplementary Table S3.

Group comparisons were performed for: 1) 0 sites vs. 1 site, 2) 0 sites vs. >2 sites, and 3) 1 site vs. >2 sites. Effect sizes were quantified as the log_10_ fold-change of group medians, such that positive values indicate higher expression or TE in the group with more ψ sites. To account for variability in site-level confidence, effect sizes were estimated using weighted regression, with weights corresponding to the number of transcripts contributing to each group. Separate effect sites were computed for protein abundance, mRNA TPM, and TE.

Statistical significance of differences between groups was assessed using the Mann–Whitney U test. Multiple testing correction was performed using the Benjamini–Hochberg procedure to control the false discovery rate.

### Statistical analysis

Experiments were performed in multiple independent experiments, as indicated in figure legends. All statistics and tests are described fully in the text or figure legend.

## Results

### Comparative analysis of pseudouridine mRNA modifications from six human cell lines using DRS

To quantify transcriptome-wide differences in psi expression across human cell types, we isolated and sequenced poly-A selected RNA from six immortalized human cell lines: i) A549, alveolar basal epithelial carcinoma; ii) HeLa, cervical carcinoma; iii) HepG2, hepatocellular carcinoma; iv) Jurkat, T-cell leukemia; v) NTERA-2, human embryonic carcinoma derived from testicular cancer; and vi) SH-SY5Y, neuroblastoma (**Fig. 1a**). The primary alignments for each sample yielded the following number of reads: 2,362,999 (A549); 4,257,869 (HeLa); 2,407,221 (HepG2); 10,931,896 (Jurkat); 10,107,973 (NTERA); 6,064,224 (SH-SY5Y; **Supplementary Table S4**. The higher number of reads observed in NTERA and Jurkat is attributed to the use of the PromethION for sequencing one or all the replicates for these cell lines. Following basecalling and genomic alignment, we mapped putative psi positions across the transcriptome for each cell line using our previously described Mod-*p* ID pipeline(16), which compares our direct RNA libraries to a reference *in vitro* transcribed (IVT) library(16, 27). An IVT control derived from the same cell type (i.e., paired-IVT or p-IVT) is preferential for this analysis; however, we have previously demonstrated that a pan-IVT comprising merged unmodified transcriptomes of different cell lines can enrich data sets where paired IVTs do not have sufficient coverage(27). We used stringent filtering criteria to reduce the possibility of false positives in psi identification: 1) IVT control has ≥ 10 reads, 2) IVT error at the U site is ≤ 10%, 3) If the paired IVT dataset has <10 reads, we used the pan-IVT dataset. For all sites that passed these criteria, we applied Mod-*p* ID to calculate *p*-values in the six cell lines (see Methods; **Figure 1b**).

**Figure 1.**
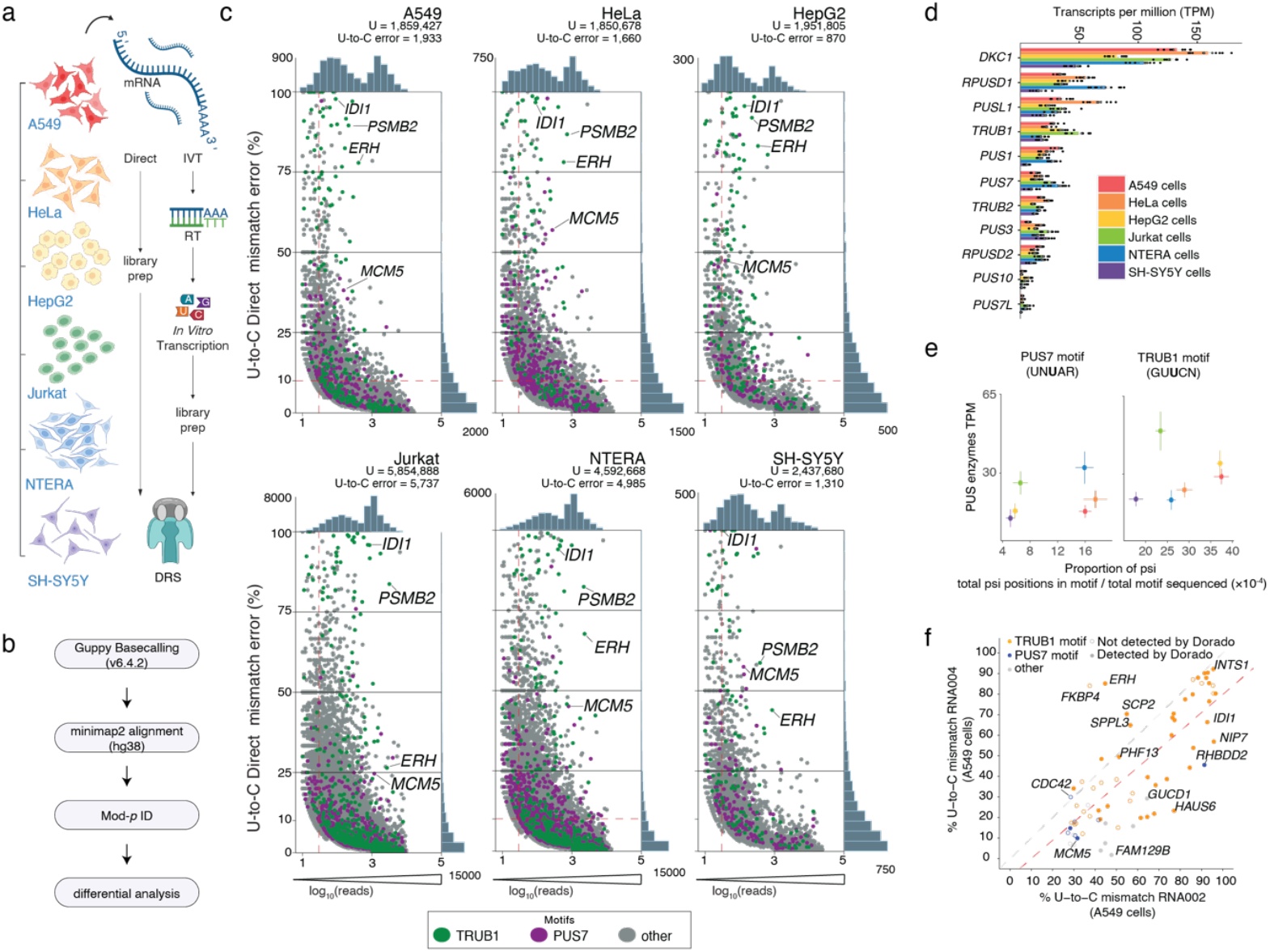
Individual cell line putative psi identification by Mod-*p* ID and PUS expression analysis. **a)** Experimental summary of A549, HepG2, HeLa, Jurkat, NTERA, and SH-SY5Y direct and IVT library prep and DRS. **b)** Overview of computational pipeline following DRS. **c)** Number of total direct reads versus the relative occupancy of putative (% mm) psi positions detected by Mod-*p* ID (*p* < 0.001) for each cell line. Grey points do not have an identified motif; colors indicate motifs for *PUS7* (purple) and *TRUB1* (green). Red dashed lines indicate baseline requirements for a position included in downstream analysis. **d)** TPMs of PUS enzymes. Individual points are from the computational resampling of the dataset. Significance from (\*\*\**p*<0.001) one-way ANOVA. **e)** (top) Fraction of the total number of reads detected by Mod-*p* ID in a *TRUB1* or *PUS7* motif divided by the total number of *TRUB1* or *PUS7* k-mer motifs observed versus the TPMs of the PUS enzyme and standard error of the mean from computational resampling. (bottom) Fraction of the total number of reads detected by Mod-*p* ID in a *TRUB1* or *PUS7* k-mer motif divided by the total number of *TRUB1* or *PUS7* k-mer motifs observed versus the relative protein expression levels of the PUS enzyme (in arbitrary units; A.U.). **f)** Scatter plot of U→C mismatch rates for 74 ground-truth Ψ sites: RNA002 (x-axis) versus RNA004 (y-axis). Filled circles = sites detected by Dorado; open circles = false negatives. Colors indicate motif context: TRUB1 (orange), PUS7 (blue), unassigned (gray).

Our first requirement for the direct library was that the putative site must have a significant difference in U-to-C basecalling error between the direct and IVT sample (*p* < 0.001). Next, we required the number of reads from the direct DRS library to be ≥ 10 for a given site. Finally, we required the U-to-C mismatch error in the direct library to be ≥ 10%. We identified putative psi sites for each cell line (**Figure 1c; Supplementary Table S5**). To find the relative ratio of putative psi sites to total uridines for a cell line, we calculated the number of putative psi sites detected by Mod-*p* ID and normalized by the number of aligned genomic uridines for each cell line. The values obtained from this normalization are 10.40×10^−4^ for A549 cells, 8.97×10^−4^ for HeLa cells, 4.46×10^−4^ for HepG2 cells, 9.80×10^−4^ for Jurkat cells, 10.85×10^−4^ for NTERA cells, and 5.37×10^−4^ for SH-SY5Y cells. Globally, A549 and NTERA cells had the highest relative ratio of putative psi sites to total uridines across cell lines, while HepG2 and SH-SY5Y cells had the lowest (**Fig 1c**).

### mRNA expression profiling of PUS enzymes across cell types

We calculated the transcripts per million (TPM) for PUS enzyme mRNAs and compared the levels across cell lines (**Figure 1d**; **Supplementary Table S6**). We categorized putative psi positions within a known psi synthase motif to test the hypothesis that higher PUS enzyme levels can result in more psi sites in each cell type. This analysis assumes that PUS mRNA levels across cell types are commensurate with their corresponding PUS enzyme levels. For the psi synthase *TRUB1*, we searched for the motif GUUCN(31); for *PUS7*, we searched for the motif UNUAR(32). We computed the proportion of Ψ positions with the motif divided by the total number of positions with the motif expressed in the transcriptome and deduced from gene models of the genome,

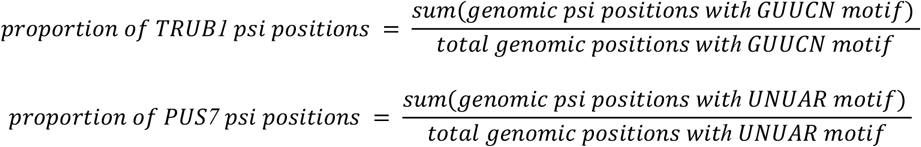

and compared these proportions to the normalized RNA expression levels for *TRUB1* and *PUS7*. Interestingly, we found that NTERA cells have the highest proportion of psi sites within a PUS7 motif (**Figure 1e**), while SH-SY5Y cells have the lowest proportion of psi for both PUS7 and TRUB1 (**Figure 1e**). These findings are consistent with our high, observed global psi levels in NTERA cells and low levels in SH-SY5Y cells (**Figure 1c**). We also found that Jurkat cells have the highest proportion of psi within a TRUB1 motif, while NTERA cells have the lowest. When comparing the proportion of psi within each of these motifs to the respective enzyme mRNA expression levels, we found no correlation between PUS mRNA expression levels in each cell type and the total number of psi sites for that cell type (**Figure 1e**). To test PUS protein expression levels we performed immunofluorescence assays, staining each cell line with fluorescently labeled Pus7 or Trub1 antibodies. We found no correlation between PUS protein expression levels and the total number of psi sites for each cell line (**Figure 1e**).

To further evaluate the robustness of ψ site detection, we leveraged improvements in nanopore chemistry introduced during the review process. Oxford Nanopore Technologies recently released a new direct RNA kit (RNA004) and basecaller, providing an opportunity to assess whether changes in sequencing chemistry affect ψ calling. Using A549 cells as a reference, we re-analyzed data with Mod-p ID on a benchmark set of orthogonally validated ψ sites (“ground truth” ψ positions) originally identified from RNA002 A549 data and confirmed by CeU-Seq(9), BID-seq(8), PRAISE seq(13), or RBS-Seq(10). The updated RNA004 chemistry produced nearly identical modification profiles, reducing U→C error frequencies below 10% at only five ψ positions (**Figure 1f**). These results demonstrate that ψ detection by Mod-*p* ID is highly reproducible and largely insensitive to changes in nanopore chemistry, supporting the robustness of our comparative ψ mapping across cell types.

### Comparison of conserved psi sites across six human cell lines

To compare the occupancy of individual psi positions across cell types, we first selected sites for which a psi modification is conserved (i.e., has been identified in every cell line) by Mod-*p* ID (*p* < 0.001). We then filtered these positions only to include those with at least 30 reads from the direct library and a minimum of 10 IVT reads for all six cell lines. This filtration step, which ensured that the mRNAs were highly expressed across all cell types, produced a list of 70 psi positions (**Supplementary Table S4**).

Of these 70 sites, we applied additional filtering criteria, including a minimum of 30% U-to-C error in each cell line, to compile a final list of “housekeeping” psi sites (**Supplementary Table S7**), used here to denote psi sites that are abundant across all cell types. Cross-validation of our 20 detected “housekeeping” sites with other methods (i.e., siRNA knockdown of a *TRUB1* or *PUS*(7, 26), CMC-mediated Illumina sequencing methods, and RNA bisulfite labeling methods(8–10, 13, 33) produced a list of 17 orthogonally validated and conserved psi sites (**Figure 2a; Supplementary Table S5**). Of these, we mapped 11 sites to the CDS and 6 to the 3’ UTR (**Supplementary Figure S2, Supplementary Table S8**). Interestingly, the *TRUB1* motif (GUUCN) predominantly appeared in 14 of our 17 highly conserved sites, and psi modification at those sites was additionally validated by a *TRUB1* siRNA knockdown experiment (7). These sites are found within genes that perform essential functions across tissue types. However, a few common pathways were identified. *NIP7, DKC1*, and *PARP4* are involved in nucleic acid processing and maintenance, and *SLC30A5* and *SLC2A1* play a role in transport functions. Interestingly, the only PUS7 substrate on this “conserved” list was *RHBDD2*, which encodes for an intramembrane serine protease necessary for processing various proteins and maintaining cellular homeostasis.

**Figure 2.**
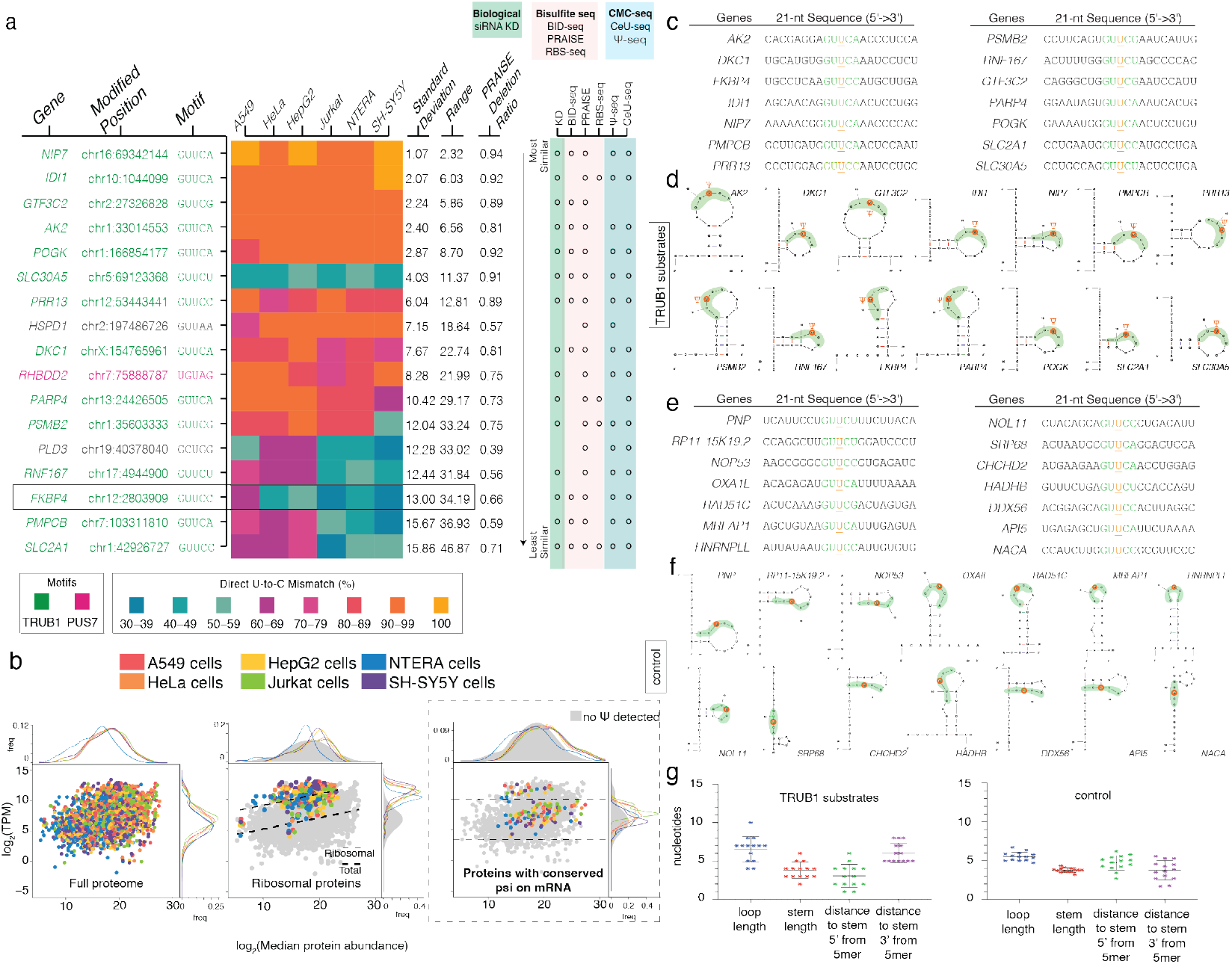
Conserved psi sites across 6 human cell lines. **a**) Heatmap of relative psi occupancy for sites identified by Mod-p ID^17^ with ≥30 reads and ≥30% U-to-C mismatch for each cell line. Colors indicate that the psi site has a TRUB1 (green) or PUS7 (pink) motif. Standard deviation and range are reported across cell types. For the same position, the PRAISE(13) deletion ratio is reported. (right) Orthogonal methods confirming the psi site including biological (green), bisulfite based (pink) and CMC-based detection (blue). **b**) Distribution of transcripts (in TPMs) with their corresponding median protein abundance for the 6 cell types (left). mRNAs encoding ribosomal proteins including those without psi on them are plotted with their corresponding protein expression (middle). The TPM of transcripts with no modification detected in any of the 6 cell lines was plotted against their median protein abundance in grey (middle). The TPM of transcripts with modifications detected in all 6 cell lines was plotted against log_2_ median protein abundance and overlaid onto the unmodified population (shown in colors for each cell line; right). **c**) 21-nt sequence of the conserved TRUB1 psi sites. **d**) Predictive secondary structure modeling of canonical 21-nt sequences of the conserved TRUB1 psi sites. **e**) 21-nt sequence of randomly selected unmodified uridine positions **f**) Predictive secondary structure modeling of canonical 21-nt sequences of the unmodified uridines **g**) Comparison of sequence properties between 14 TRUB1 substrates and 14 control sequences. The bars show the median value with standard error.

Next, we calculated the standard deviation (SD) for each psi position to compare across all six cell lines (**Figure 2a**). The conserved psi site located on *NIP7* (chr16:69342144) has the smallest SD positional occupancy ranging from 98% (NTERA) to 100% (A549, HepG2, and SH-SY5Y). *NIP7* (Nucleolar pre-rRNA processing protein) is involved in RNA binding activity for ribosomal biogenesis. We also observed in a previous study that this position on *NIP7* is relatively static in SH-SY5Y cells across multiple treatments(7). Conversely, the relative positional occupancy of the psi on *SLC2A1* (chr1:42926727) ranges from 32% U-to-C error (Jurkat) to 79% (HepG2). *SLC2A1* (solute carrier family 2 member 1) is a critical glucose transporter essential for tissue-specific energy delivery. While *SLC2A1* is present in both liver and T cells, its primary role in the liver is related to supporting glucose metabolism, whereas in T cells, *SLC2A1* is crucial for immune function and response. This psi site has also been found to have stable occupancy across different treatments for SH-SY5Y cells(7).

### Relationship between transcript levels and positional psi occupancy

The conserved transcript *FKBP4* (chr12:2803909) has highly variable psi occupancy across the cell lines ranging from 38% (Jurkat cells) to 69% (A549 cells), as well as high variability in transcript expression levels ranging from 81±8 (SH-SY5Y cells) to 735±16 (NTERA cells) TPMs (**Figure 2a; Supplementary Figure S2**). *FKBP4* (prolyl isomerase 4) stabilizes and manages steroid hormone receptors in both lung and T cells. FKBP4 affects inflammation and immune responses in the lung, while FKBP4 regulates cell differentiation and effector functions in T cells.

We observed several examples of an inverse relationship between mRNA levels and psi positional occupancy. In NTERA cells, *FKBP4* expression is the highest out of all the cell lines (349 TPMs) but one of the lowest psi positional occupancies at 42%. Conversely, A549 cells have significantly lower *FKBP4* expression levels with 111±6 TPMs but show the highest psi positional occupancy at 69%. *SLC2A1* is also differentially expressed with variable psi occupancy at chr1:42926727 across the cell lines. However, this case directly correlates transcript abundance and psi positional occupancy. HepG2 cells have the highest expression of *SLC2A1* at 210±9 TPMs and the highest U-to-C mismatch at 79%, and Jurkat cells have the lowest expression and psi occupancy at 37±4 and 32%, respectively (**Figure 2a; Supplementary Figure S2**).

### Protein expression of transcripts with conserved psi sites

To assess post-transcriptional regulation by psi, we performed a joint mRNA and protein analysis for the conserved positions. This was an ideal group for which to perform this analysis because these positions were found within highly expressed targets, and the levels of psi (based on U-to-C error) were high. We performed quantitative mass spectrometry on all 6 cell lines, labeling each with a unique isobaric tandem-mass-tag (TMT) (see Methods; **Supplementary Table S2**).

We first analyzed all measured transcripts, agnostic to the presence of psi, to visualize the full proteome distribution for each of the 6 cell lines (**Figure 2b; Supplementary Table S3**). Overall, the mRNA levels are weakly correlated with corresponding protein abundance. Additionally, the genes represented by NTERA cells had lower corresponding protein expression than the other 5 cell lines.

Next, to assess post-transcriptional regulation by psi we applied several stringent criteria to qualify for analysis: at least one genomic uridine position on a given transcript must have 30 reads and contain a conserved psi site with a minimum of 10% direct U-to-C mismatch in all six cell lines. Filtering with these criteria resulted in 68 unique transcripts. We further filtered these targets to include only those with a protein reading in each of three replicates resulting in 22 targets, with psi found primarily in the CDS (**Figure 2b, Supplementary Figure S3, Supplementary Table S3**). Interestingly, ∼68% of the 68 psi-bearing mRNA targets did not have protein detected despite high levels of mRNA expression (>30 TPMs) compared to 87% of non-psi-bearing transcripts that did not have protein detected despite high levels of mRNA expression (>30 TPMs). This suggests that psi-bearing mRNA targets may have higher protein expression levels because they are more frequently exceeding the minimum threshold of detection. To confirm this observation, we compared the relative lengths of the proteins detected and found that the median protein lengths of the modified and unmodified groups are similar. We then ascertained whether 32% (proteins found in the psi-bearing group) is statistically different from 13% (proteins found in the non-psi bearing group) by performing a simulation in which 68 genes were randomly sampled from a dataset containing gene names of mRNA transcripts with more >30 reads. A total of 10 million randomly selected sets of 68 were generated and cross referenced to a dataset containing genes for which proteins were identified via mass spectrometry in all 3 replicates of the 6 cell lines. The 95% confidence interval for the mean and standard deviation of this distribution is [13.869;13.874] and [4.148; 4.153] which suggests that the empirically observed difference of 32% and 13% between the modified and unmodified populations was not due to sample size imbalance and supports the observation that pseudouridylation impacts protein expression.

Consistent with this observation, we noted an increase in protein expression for psi-bearing transcripts of similar expression levels compared to the protein expression of non-psi-bearing transcripts. This is visualized by the average position of psi transcripts on the y-axis (TPMs) in the scatter plot of **Figure 2b** (right), yet higher-than-median protein count values along the x-axis of the same scatter plot. One exception to this pattern is with NTERA cells that is left-shifted compared to the rest of the population, indicating lower than median protein abundance for these targets. To account for the sample size imbalance between the modified (68) and unmodified (3400) transcripts, a down sampling simulation was performed. Both datasets were pooled and 10 million sets of 68 randomly sampled genes were generated. Each random set of 68 genes was subsequently cross referenced to a comprehensive dataset containing all transcripts with protein detected in all 3 replicates of 6 cell lines, irrespective of modification status. The portion of transcripts for which a protein was detected in each set was then calculated. The resulting 95% confidence interval for the mean and standard deviation of this distribution is [13.871; 13.877] and [4.148; 4.153], respectively. This suggests that the empirically observed 32% of proteins belonging to modified transcripts which met the detection threshold, compared to 13% for unmodified transcripts, is not due to random chance and the difference between the two groups is statistically significant.

We also observed two separate populations for transcripts with a conserved psi, specifically a population of targets with high mRNA levels and relatively lower protein expression than expected based on the mRNA levels (**Figure 2b**, right). We observed that the targets in the top population were mainly encoding ribosomal proteins. To test the hypothesis that mRNAs encoding ribosomal proteins typically have high mRNA expression level yet lower protein counts, we plotted all mRNAs encoding ribosomal proteins including those without psi on them (**Figure 2b**, middle) and observed a tight population that has a significantly higher trendline than the global correlation trendline for all mRNAs.

### Secondary structure modeling of the region surrounding psi position for TRUB1 targets

As mentioned earlier, most highly conserved targets were TRUB1 substrates (**Figure 2a**). Previous studies have analyzed the structural features standard to TRUB1 targets in a single cell line(31) and found that the stem and loop structure on mRNA within the TRUB1 motif is sufficient for mRNA pseudouridylation. Thus, we aimed to assess whether these structural features were present on sites where psi is conserved across cell lines. Using Mfold(34), we took a 21-mer sequence flanking the conserved and validated psi positions within a TRUB1 motif (14 targets) and predicted the RNA secondary structure (**Figure 2c,d; Supplementary Table S9**). The 21-mer sequence was chosen because the predictions are more accurate with shorter sequences(34). Additionally, we randomly selected 14 control sites for each target, a total of 196 sites with the same motif but bearing no psi modification (**Figure 2e, f; Supplementary Table S9**).

In comparing the structural characteristics of these groups, we observed that the control sites exhibit shorter loop lengths on average compared to the psi sites (p = 0.0369 < 0.05). While the stem lengths were not significantly different between control sites and psi sites (p = 0.9723), the 5mer motif for psi sites tends to be closer to the 5’ end of the stem than in the control sites (p = 0.0016 < 0.05). Conversely, the 5mer motif for controls is closer to the 3’ end of the stem than psi targets, and this difference is significant (p < 0.0001; **Figure 2g**).

We randomly selected 14 control sites from the 196 total controls to directly compare with the 14 psi targets (**Figure 2d,f)**. Interestingly, some control sites displayed structural variations distinct from those observed in psi targets. For instance, SPR68 exhibits its 5-mer motif beyond the 5’ end, while NACA positions its first nucleotide of the 5-mer at the 3’ end of the stem, with four nucleotides extending behind the stem. In contrast, all psi targets exhibit the 5-mer motif positioned within the stem, loop, or both.

Importantly, all psi targets displayed their U/psi positioned within the loop of hairpin structures for the 21-mer structures, whereas only 7 of the 14 control sites had U/psi positioned within the loop. Additionally, 3 of the 14 control sites contained internal loops, a feature not observed in psi targets. We also compared the U/psi position within 31-mer and 41-mer structures for the 14 psi targets and 14 controls. For psi targets, 12 out of 14 have their psi sites on the loop for 31-mer and 41-mer structures (**Figure 2d**; **Supplementary Table S9, Supplementary Figure S4-5**). Therefore, longer sequences do not significantly change the preference for loop locating of psi sites. As for the U controls, 3 U/psi sites are located on the loop for the 31-mer structures and 4 for the 41-mer structures. Both 31-mer and 41-mer structures have 8 U/psi sites located on the stem. In summary, for all lengths of structures, U/psi sites prefer to be located on the loop of psi targets. Conversely, the control sites appear in the loop less often.

### Site-specific psi presence or absence can be cell-type dependent

The unique presence or absence of a modification site in a cell type implies some form of epitranscriptome-level specialization for that cell type. We were interested to see whether psi sites were present only in specific cell lines, even if RNA expression levels were conserved. We constructed a table of Mod-*p* ID detected and orthogonally confirmed psi sites where the modification was detected in 3 of the 6 cell lines (**Figure 3a; Supplementary Table S10**). For example, *DAZAP1* (chr19:1434893) was only present in A549, HeLa, and HepG2 while *ITFG1 (*chr16:47311407) was present only in A549, HepG2 and SH-S75Y cells. Interestingly, for this group of modified sites, we found no positions within this category that were hypermodification type 1(16), meaning all identified sites had <40% U-to-C mismatch, indicating <50% psi occupancy. We compared all aligned reads’ raw ion current traces at the psi position, revealing that psi presence affects the ion current distributions (**Figure 3b**).

**Figure 3.**
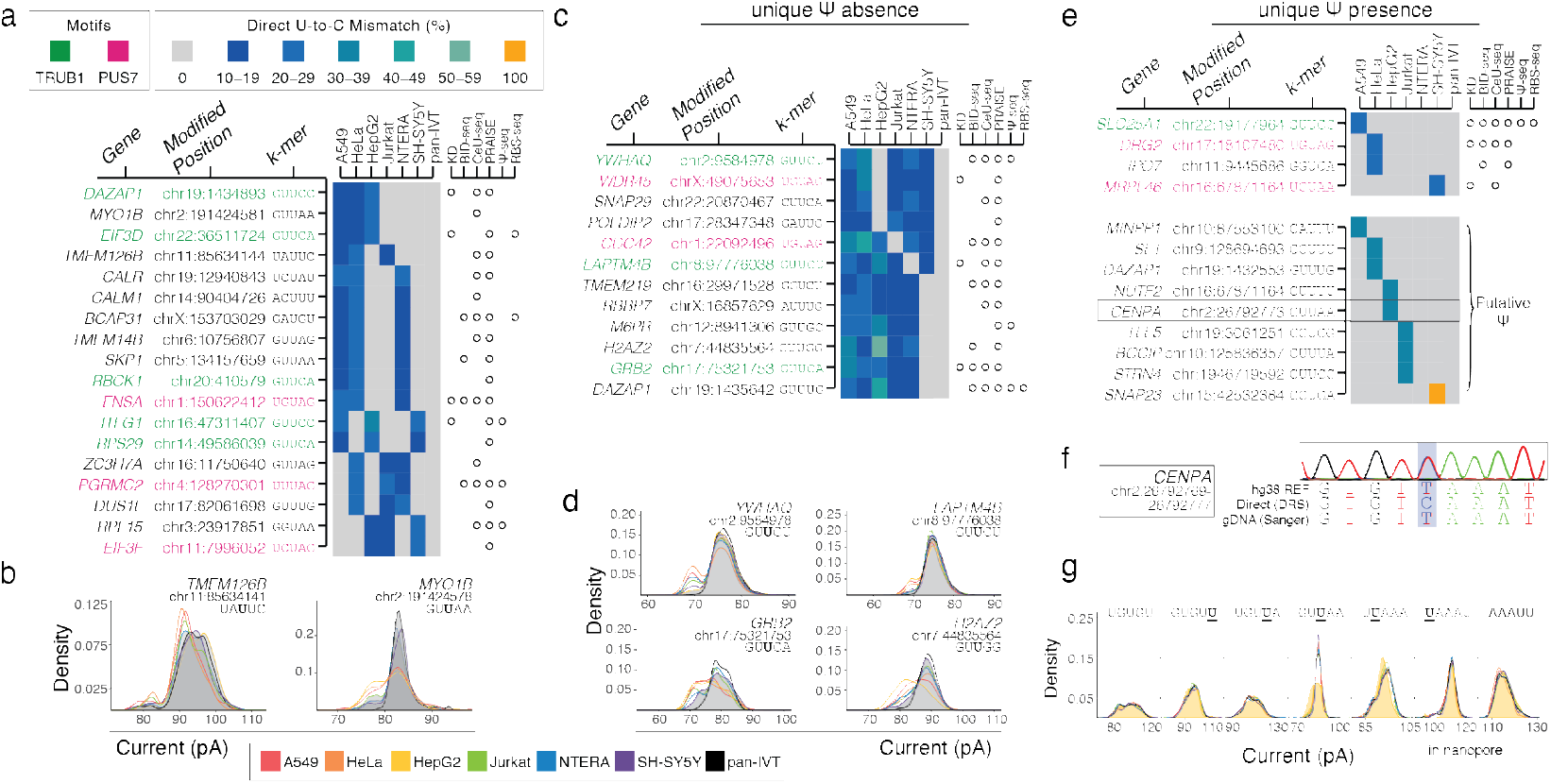
Cell type-specific expression of psi. **a**) Heatmap of relative psi occupancy for sites detected in 3 cell lines identified by Mod-*p* ID and confirmed by orthogonal methods. Colors indicate that psi site has a TRUB1 (green) or PUS7 (pink) motif. **b**) KDE ionic current traces of cell-type split presence/absence of psi site. Colors indicate traces for each cell line. The cell line with no psi detected is a grey-shaded distribution. **c**) Heatmap of the unique absence of a cell line identified by Mod-p ID and confirmed by orthogonal methods. Colors indicate that psi site has a TRUB1 (green) or PUS7 (pink) motif. **d**) KDE ionic current traces of cell-type specific absence of psi site. Colors indicate traces for each cell line. The cell line with no psi detected is a grey-shaded distribution. **e**) Heatmap of cell type-specific unique presence of putative psi sites’ relative occupancy. **f**) Sanger sequencing of cell type-specific putative psi positions *CENPA* **g**) KDE ionic current traces of *CENPA* for +/-3 positions. Colors indicate traces for each cell line. The cell line detected to have the putative psi is shaded with its corresponding color. pan-IVT trace is in black.

The unique absence of a psi site for a single cell line may implicate cell type-specific functionality. To study this, we had to change our selection criteria from what was used to analyze “housekeeping” psi targets (i.e., 30 direct reads and >30% U-to-C error in every cell line). We relaxed the criteria to include positions with at least 30 direct reads in every cell line and >10% psi positional occupancy for the detected site in 5 out of 6 cell lines (**Supplementary Table S11**). We detected 38 putative psi sites with a unique absence in a single cell line with Mod-*p* ID, 12 of which are orthogonally confirmed psi positions (**Figure 3c**). All 12 sites are present in A549 and HeLa cells, indicating no unique absence of psi at any detected position in these two cell lines. SH-SY5Y demonstrates the highest occurrence of uniquely absent sites, with only 6 psi sites identified. Four of the six psi positions present in SH-SY5Y are found on transcripts (*CDC42* (chr1:22092496), *WDR45* (chrx:49075653), *YWHAQ* (chr2:9584978), and *SNAP29* (chr22:20870467)) related to neuron function directly in the cellular activities or through processes crucial for neuronal health and activity. Each of these 12 positions was orthogonally validated to be psi (**Figure 3c**). The position with the largest difference between the two cell lines was within *H2AZ2* (chr7:448355564), which is hypermodified (type I) in HepG2 cells and not modified in SH-SY5Y cells while being highly expressed in both. This gene encodes a variant of the histone H2A complex, plays a role in synaptic plasticity in neurons, and plays a role in liver development. We do not know which PUS enzyme is responsible for this modification. However, this position was confirmed to be psi by two orthogonal methods(8, 13).

To further confirm the cell-type-specific absence of psi, we overlaid the per-read ionic current intensity distribution ionic current traces for all aligned reads to the psi position (**Figure 3d**). As expected, at chr2:9584978 (*YWHAQ*), we observed a single, smooth distribution in HepG2 cells and the pan-IVT, whereas, in the other five cell lines, there is an additional peak to the left of the expected distribution peak, indicating a difference in the physical properties of the sequences (i.e., thus leading to a change in the ionic current disruption) corresponding to the observed absence of psi (**Figure 3d**).

Likewise, the unique presence of psi positions in a cell line may further provide information on the cell-type-specific functionality of psi. Of the 63 orthogonally confirmed psi sites using Mod-*p* ID present in only one cell line with at least 10% relative positional occupancy, 7 sites have a *TRUB1* motif, and 12 have a *PUS7* motif. Only 4 of the orthogonally confirmed psi sites have above a 20% U-to-C error signature: *DRG2* (chr17:18107480), *MRPL46* (chr14: 88467298), *SLC25A1* (chr22:19177964), and *IPO7* (chr11:9445686). Additionally, we identified 14 putative psi sites using Mod-*p* ID which have >30% U-to-C errors (**Figure 3e**). For all sites of unique modification presence, we ruled out the presence of a single-nucleotide variant in these cell lines by Sanger sequencing on the corresponding gDNA (**Figure 3f; Supplementary Figure S4**). We wanted to confirm that these sites deviated from the expected ionic current, which indicates a probable modification. We observed that the pan-IVT trace and the other five cell lines have similar ionic current trace distributions and a visible deviation of the cell line distribution with the modification (**Figure 3g**).

### Type II hypermodification of transcripts can be variable or shared across cell lines

Type II psi hypermodification has been previously defined as transcripts with more than one psi modification(16). To assess the type II hypermodification status across cell types, we filtered to require ten direct reads, and the paired-IVT had ten reads at the query position (**Supplementary Table S12**). We detected up to 5 modified positions per transcript for A549 cells, HeLa cells, HepG2 cells, Jurkat cells, and NTERA cells. We detected the presence of many transcripts with two modifications in all cell lines. A549 cells, NTERA cells, and Jurkat cells consistently had the highest detected type II hypermodified transcripts. In contrast, SH-SY5Y cells had the lowest number of hypermodified transcripts as we only detected up to 3 positions on one transcript (**Figure 4a**).

**Figure 4.**
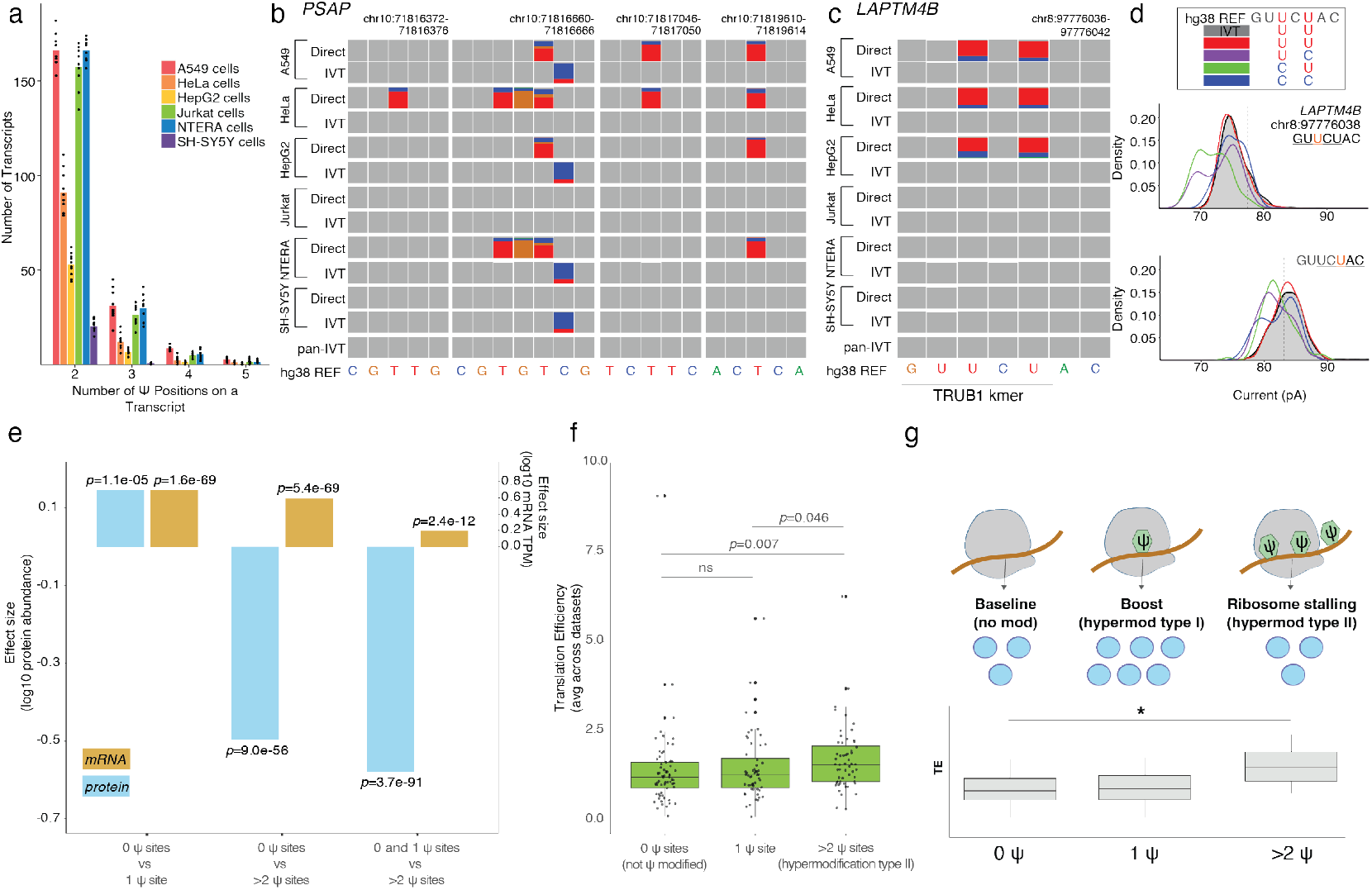
Type II hypermodification analysis. **a)** Number of hypermodified type II transcripts with 2, 3, 4, and 5 putative psi sites. **b)** Variable type II hypermodification across cell lines for the prosaposin gene, *PSAP*. Location on the transcript of three putative psi sites and IGV snapshots of each position’s motif for the six-cell line’s direct and IVT library and pan-IVT. **c)** Special case of type II hypermodification: double modification where multiple putative psi positions are detected within a motif on *LAPTM4B*. IGV snapshot of each position’s motif for the six-cell line’s direct and IVT library and pan-IVT. The first detected modification is found in *TRUB1* motif. **d)** Merged cell line KDE ionic current traces of the two double mod putative psi sites for all modification combinations for the two sites in different colors. The ionic trace for the pan-IVT is shaded in grey. **e)** Bar plot shows the weighted regression coefficient (effect size) of protein abundance and mRNA expression in 3 comparison groups: mRNAs with 0 vs 1 ψ-sites, 0 vs >2 ψ-sites, 0 and 1 vs >2 ψ-sites. Protein abundance was determined by mass spectrometry. mRNA expression was measured in TPMs from nanopore DRS results. Statistical analysis was conducted using the Mann-Whitney U test, p-values are reported for each comparison group. **f)** Translation efficiency (TE) values were calculated from publicly available ribosome profiling (Ribo-seq) and RNA-seq datasets obtained from GEO. For each transcript, TE was defined as the ratio of ribosome-protected fragment counts to RNA seq counts. Statistical analysis was conducted using the Kruskal-Wallis test, p-values are annotated for each comparison group. **g)** Model showing the ribosome (grey) interacting with mRNA (brown line) in the absence and presence of ψ (green). Protein output is shown with blue circles and as a function of no modification, hypermodification type I and hypermodification type II. Statistical analysis was conducted using the Kruskal-Wallis test (*p<0.05).

*PSAP* encodes the prosaposin gene and is one example of a transcript with multiple positions orthogonally confirmed (chr10:71816374(10, 13), 71816664(8–10, 13, 33), 71817048(13), and 71819612(13)) and detected at varying positional occupancy with our Mod-*p* ID method (**Figure 4b**). The encoded protein is essential for the proper functioning of the lysosome. Of the positions identified, chr10:71819612 is in the CDS, while the other three are in the 3’ UTR. None of these confirmed sites have a known PUS motif. All 4 psi locations are present in HeLa cells, 3 in A549 cells, 2 in NTERA, 1 in HepG2, and 0 in Jurkat and SH-SY5Y. The positional occupancies for detected psi sites are relatively low, with a maximum of 33% in HepG2 at chr10:71819612. Mod-*p* ID also identified a 5^th^ putative position at chr10:71816662 in HeLa and NTERA cells. This additional putative psi site falls within the same motif of chr10:71816664, a putative double modification (**Figure 4b**).

*LAPTM4B* (chr8:97776038, 97776040) is another example of a double modification, where chr8:97776038 has a TRUB1 motif, GUUCU (**Figure 4c**), which has been orthogonally confirmed with both PRAISE(13) and CeU-seq(9). We confirmed with Sanger sequencing that neither position was an SNV (**Supplementary Figure S4**). CMC and bisulfite-based methods for psi detection cannot detect two neighboring modifications. Thus, there are no known examples of double modifications. Double modifications can also potentially disrupt current traces in direct RNA sequencing. To explore further, we performed a strand analysis of all six cell lines’ combined reads (pan-direct) to see if the double modification occurred simultaneously in both positions on a given read. A read must span both positions and only contain a U or C to be included in the analysis. We separated the 1,823 reads into four groups: i) no modification (1,619 reads), ii) U-to-C error in the first position and an unmodified second position (76 reads), iii) an unmodified first position and U-to-C error in the second position (124 reads), and iv) double modification (4 reads). We wanted to visualize the ionic current distribution at these two positions, specifically when the nucleotide of interest is in the middle position of the nanopore, for each grouping. We observe changes in the distribution pattern corresponding to each grouping. Overall, this suggests that singly modified and doubly modified populations may exist within the same motif, but the mechanism remains unknown (**Figure 4d**).

### Type II hypermodifications correlates with elevated translation efficiency but reduces protein abundance

We further assessed if the presence of ψ modifications and the number of sites on a transcript had an effect on expression and translation efficiency. Our analysis (see **Methods**) show that while ψ presence consistently correlates with higher transcript abundance, protein output follows a biphasic pattern: transcripts with a single ψ site exhibit elevated protein levels, whereas those with two or more ψ sites show reduced protein abundance (**Figure 4e**). Strikingly, translation efficiency (TE) remains high for transcripts containing multiple ψ sites (**Figure 4f; Supplementary Table S3**). This paradox is consistent with a ribosome stalling model in which hypermodified mRNAs (high-occupancy sites; U-to-C error >40%) recruit ribosomes more efficiently but experience increased pausing, leading to reduced protein yield despite elevated TE (**Figure 4g**).

## Discussion

Our recently developed method, Mod-*p* ID(16), enables transcriptome-wide mapping of pseudouridine (ψ) sites and quantitative comparison of psi occupancy levels across conditions. Multiple orthogonal techniques have been developed to detect psi at single-nucleotide resolution, each with distinct strengths and limitations. Mod-*p* ID achieves comparable levels of de novo site discovery to these established methods, identifying roughly 44 % of sites not validated by other approaches—within the 55–60 % unvalidated range reported for alternative methods (**Supplementary Figure S6**). This uncertainty is inherent to all transcriptome-wide detection strategies, as chemical derivatization, amplification bias, and alignment artifacts can each obscure a fraction of true modifications. In previous work, psi sites uniquely detected by Mod-*p* ID were orthogonally validated by CLAP (n = 9)(7, 35), demonstrating that nanopore-based detection can reveal bona fide sites missed by short-read sequencing.

The primary advantage of Mod-*p* ID is not increased detection yield but expanded analytical scope. By leveraging long-read direct RNA sequencing, Mod-p ID can detect multiple modifications on a single RNA molecule, enabling identification of hypermodified transcripts (type II(16)) that cannot be resolved by short-read or chemical-based approaches. This capability allows the examination of psi stoichiometry, positional context, and co-occurrence across the same RNA strand—features inaccessible to other transcriptome-wide ψ-mapping methods. We applied this framework to six human cell lines (A549, HeLa, HepG2, Jurkat, NTERA, and SH-SY5Y) and observed substantial variation in psi density and motif distribution among them (**Figure 1**).

Interestingly, psi enrichment patterns do not correlate with PUS7 or TRUB1 levels (**Figure 1e**), indicating that psi stoichiometry across cell types is regulated by factors beyond enzyme expression—possibly including substrate accessibility, RNA structure, or post-translational regulation of PUS activity.

To confirm the robustness of psi detection, we re-evaluated Mod-*p* ID performance using Oxford Nanopore’s updated direct RNA chemistry (RNA004) and basecaller (**Figure 1f)**. Using A549 cells as a benchmark, we compared previously validated psi sites—confirmed by CeU-Seq(9), BID-seq(8), PRAISE(13), or RBS-Seq(10)—and found that RNA004 preserved nearly all modification calls, reducing U→C error rates below 10 % at only five positions. These results confirm that psi detection by Mod-*p* ID is highly reproducible and largely unaffected by changes in nanopore chemistry, underscoring the stability of the ψ-specific current signature.

Analysis of conserved psi sites in highly expressed housekeeping mRNAs revealed that most of these positions (> 80 %) are substrates for TRUB1. Some conserved sites displayed near-complete occupancy across cell types (e.g., *NIP7*, > 97.6 %), while others showed variable occupancy (e.g., *SLC2A1*, 40–90 %), suggesting site-specific regulation of psi stoichiometry. Many of these transcripts encode proteins involved in ribosome biogenesis (e.g., *NIP7*(36), *DKC1*), consistent with psi’s proposed role in maintaining translational homeostasis. Predictive secondary-structure modeling of conserved TRUB1 psi sites confirmed enrichment within hairpin loops, consistent with prior structural analyses (**Figure 2**) (31).

We also observed cell type–specific psi sites whose occupancy varied across cell lines. Sites present in only a subset of cell types typically showed lower relative occupancy (10–50 %), possibly reflecting dynamic regulation near the detection threshold. Raw ionic current analysis confirmed distinct signal shifts even at low occupancy, supporting the presence of true modifications (**Figure 3f**). Some of these sites may underlie specialized translational programs; for example, a HeLa-specific psi site in *DRG2*, a PUS7 target, could contribute to the regulation of cell proliferation and microtubule dynamics(37).

Beyond single modifications, we identified transcripts bearing multiple closely spaced psi sites, including rare double modifications within the same motif. Such events represent a unique category of type II hypermodification, where two psi residues occupy positions simultaneously within the ∼11-nt region interacting with the nanopore. These configurations are invisible to short-read methods but detectable in long-read ionic current traces(18). For example, *LAPTM4B* harbors double psi modifications in A549, HeLa, and HepG2 cells at chr8:97776038 and chr8:97776040 (**Figure 4**). The visible differences in the KDE of merged ionic traces may indicate that the detection of double modification is not a false positive. chr8:97776038 has been orthogonally confirmed(9, 13). TRUB1 siRNA knockdown in HeLa cells showed the U-to-C error disappear in both chr8:97776038 and chr8:97776040 compared to both the native HeLa mRNA and scrambled control. We ruled out the presence of an SNV in all cell lines for these sites by Sanger sequencing on the corresponding gDNA (**Supplementary Figure S1**). These findings demonstrate that psi can occur in dense clusters on mRNAs, supporting a model in which modification “hotspots” modulate translation through additive or cooperative effects.

Functional studies of psi have shown the modification can influence translation efficiency and induce changes in the translation of the mRNA itself(5, 38, 39). Our data show that mRNAs bearing high-occupancy, conserved ψ sites produced significantly higher protein levels than unmodified transcripts, consistent with prior reports linking psi to enhanced translation efficiency (28). This trend was not universal—NTERA cells, for instance, displayed lower-than-median protein abundance for psi-modified transcripts—suggesting that psi’s translational effects depend on cell-specific regulatory contexts or isoform usage. Integrating ribosome profiling (Ribo-seq) with Mod-*p* ID data will help delineate whether psi primarily enhances elongation efficiency, affects ribosome pausing, or stabilizes transcripts under specific conditions.

Taken together, these findings move beyond incremental mapping of psi to reveal a quantitative and mechanistic framework for understanding how ψ regulates translation. By combining long read psi profiling, proteomics, and ribosome footprinting, this study provides new evidence that ψ stoichiometry and spatial organization are key determinants of translational output. Mod-p ID’s ability to detect co-occurring modifications on individual RNA molecules establishes a foundation for future efforts to model RNA modification networks at single-molecule resolution and to explore how programmable pseudouridylation could be leveraged to modulate translation in therapeutic contexts.

## Supporting information

Legends for supplementary tables

## Code Availability

All code used in this work is publicly available at https://github.com/RouhanifardLab/PanHumanPsiProfiling

## Data Availability

The data that support this study are available from the corresponding author upon reasonable request. Sequences were aligned to genome version hg38.p10. Unless otherwise stated (*See Accessing Publicly Available Data Sets*), all FASTQ files and Fast5 raw data generated in this work are publicly available in NIH NCBI SRA under the BioProject accession PRJNA1108269. Due to the size of the supporting data bam files and Mod-*p* ID outputs are hosted on Dropbox at https://www.dropbox.com/scl/fo/p939edmv71o5cwl9by5x0/AFbhroxm3LIyaGY_kzgODHM?rlkey=nd6lgzg3usahxwiq8d3k2vbj6&st=uss6f43a&dl=0 Proteomics data is available on MassIVE (MSV000095673) and the ProteomeXchange (PXD055082).

## Acknowledgments

S.H.R. and M.W. acknowledge support from NIH R01HG011087 and NIH R01HG012856 as well as support through an Opportunity Fund by the Technology Development Coordinating Center at Jackson Laboratories (NHGRI federal award no. U24HG011735). We thank the Mass Spectrometry facility associated with the Chemistry and Chemical Biology Department at Northeastern University, which provided TMT-labeling and LC-MS/MS services.

## Author Contributions

C.A.M. and S.H.R. conceived of the research. C.A.M. designed the experiments. C.A.M., Y.Q., O.F., D.B., and I.N.K. performed experiments and sequencing runs. C.A.M., Y.Q., M.T., Y.L., M.M, and O.F. analyzed the data with guidance from S.H.R., M.J. and M.W. C.A.M. and M.M. wrote the paper with guidance from S.H.R.

