## Supplementary material for "mRNA psi profiling using nanopore DRS reveals cell type-specific pseudouridylation and translational regulation": Legends for supplementary tables

**Supplementary Table S1**: Primers used for Sanger sequencing

**Supplementary Table S2**: Reporter labeling scheme and MaxQuant output-evidence table

**Supplementary Table S3**: Normalized protein abundance and corresponding transcripts per million (TPM) in 6 cell lines and translation efficiency values. Corresponding TE values for each transcript in the second tab.

**Supplementary Table S4**: Sequencing device used and read counts for each cell type

**Supplementary Table S5**: Output of Mod-p ID; Putative psi sites for each cell line, not filtered for read count cutoffs, IVT cutoffs or mismatch cutoffs. Orthogonal validation is indicated in this table for individual sites.

**Supplementary Table S6**: Transcripts per million for the mRNAs encoding psi synthase (PUS) enzymes for each cell line

**Supplementary Table S7**: Lookup table for 70 conserved psi sites

**Supplementary Table S8**: Table of psi sites and the location of the psi on the mRNA body

**Supplementary Table S9**: 21mer sequences bearing a TRUB1 motif with and without a psi position.

**Supplementary Table S10**: Table of Mod-p ID detected sites that are orthogonally validated and the modification was detected in 3 of 6 cell lines.

**Supplementary Table S11**: Psi modified versus unmodified transcripts and corresponding protein expression with statistical analysis.

**Supplementary Table S12**: Type II hypermodification analysis across cell types.
