## Supporting Information for "mRNA psi profiling using nanopore DRS reveals cell type-specific pseudouridylation and translational regulation"

**Table of Contents**

**Supplementary Figure 1**………………………………………….………….………………2

*Sanger sequencing of putative psi sites’ corresponding gDNA for SNV detection*

**Supplementary Figure 2**………………………………………….………….………………3

*Conserved psi across 6 human cell lines*

**Supplementary Figure 3**………………………………………….………….………………4

*Expanded view of mRNAs with conserve psi sites and their corresponding protein abundance*

**Supplementary Figure 4**………………………………………….………….………………5

*mFold structures of sites with psi within 21-, 31-, and 41-mer sequence context*

**Supplementary Figure 5**………………………………………….………….………………6

*mFold structures of sites without psi within 21-, 31-, and 41-mer sequence context*

**Supplementary Figure 6**………………………………………….………….………………7

*Comparison of Mod-p ID against non-nanopore orthogonal methods for pseudouridine detection*

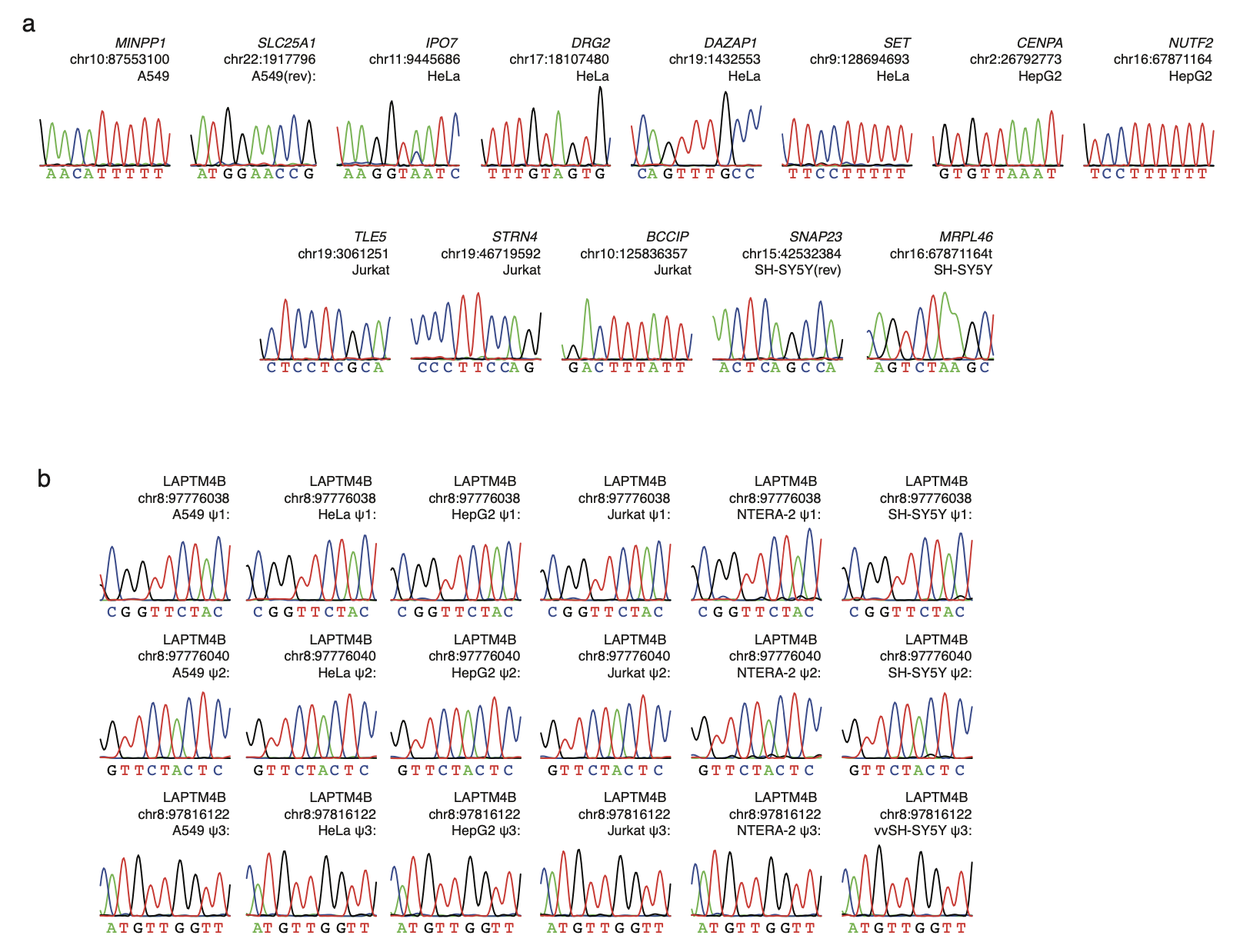

**Supplementary Figure 1.** a. Sequence chromatograph of putative, cell-type specific psi modification sites’ corresponding gDNA. The center position in the reference sequence corresponds to the putative psi. b. Sequence chromatograph of psi positions with hypermodification type II for their corresponding gDNA sequence. The center position in the reference sequence corresponds to the putative psi.

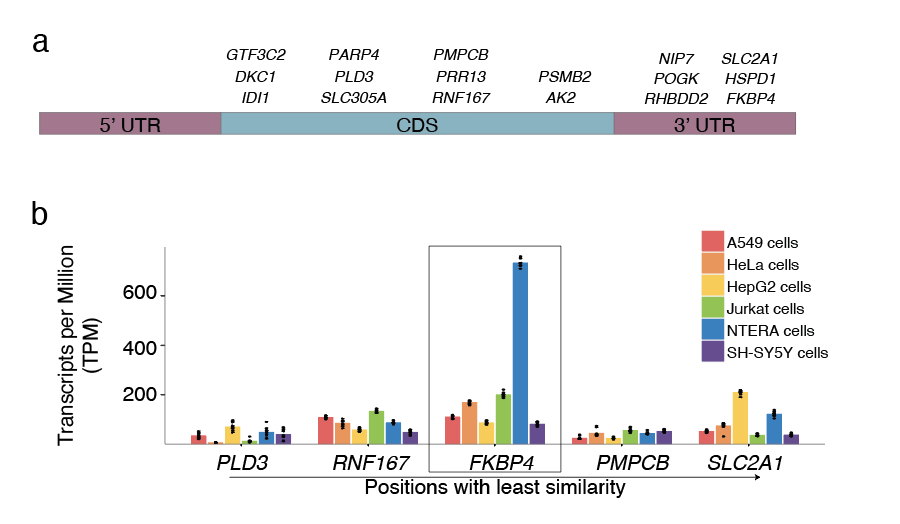

**Supplementary Figure 2.** Conserved psi sites across 6 human cell lines

**a**) Transcript location of psi sites. **b**) TPMs of transcripts bearing conserved psi positions with the least similarity for each cell line. Least similarity indicates a high standard deviation in the U-to-C error at a position within the reported transcript (i.e., bigger differences in occupancy across cell lines).

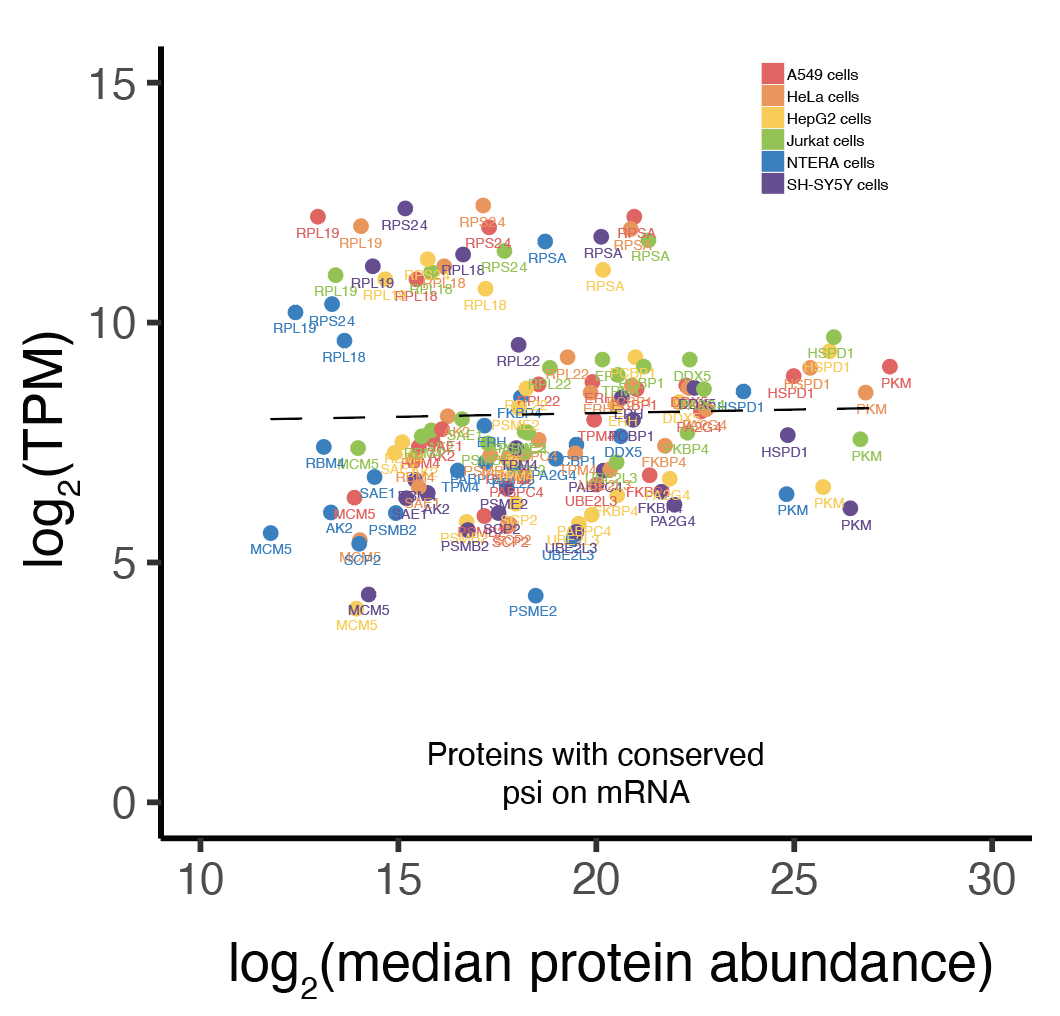

**Supplementary Figure 3.** Distribution of transcripts (in TPMs) with their corresponding median protein abundance for 6 cell types for that have a conserved psi position within them (shown in colors for each cell line). Note that this is the same as the Figure 2b panel but expanded and with labels for the individual proteins.

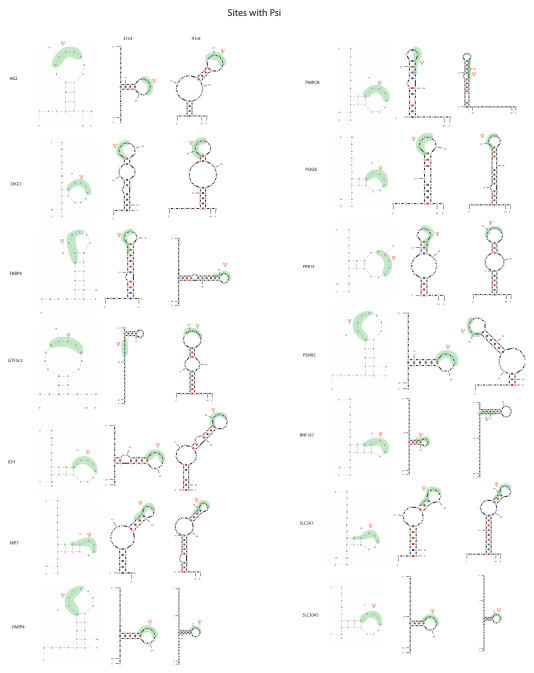

**Supplementary Figure 4.** mFold structures of sites with psi within 21-, 31-, and 41-mer sequence context

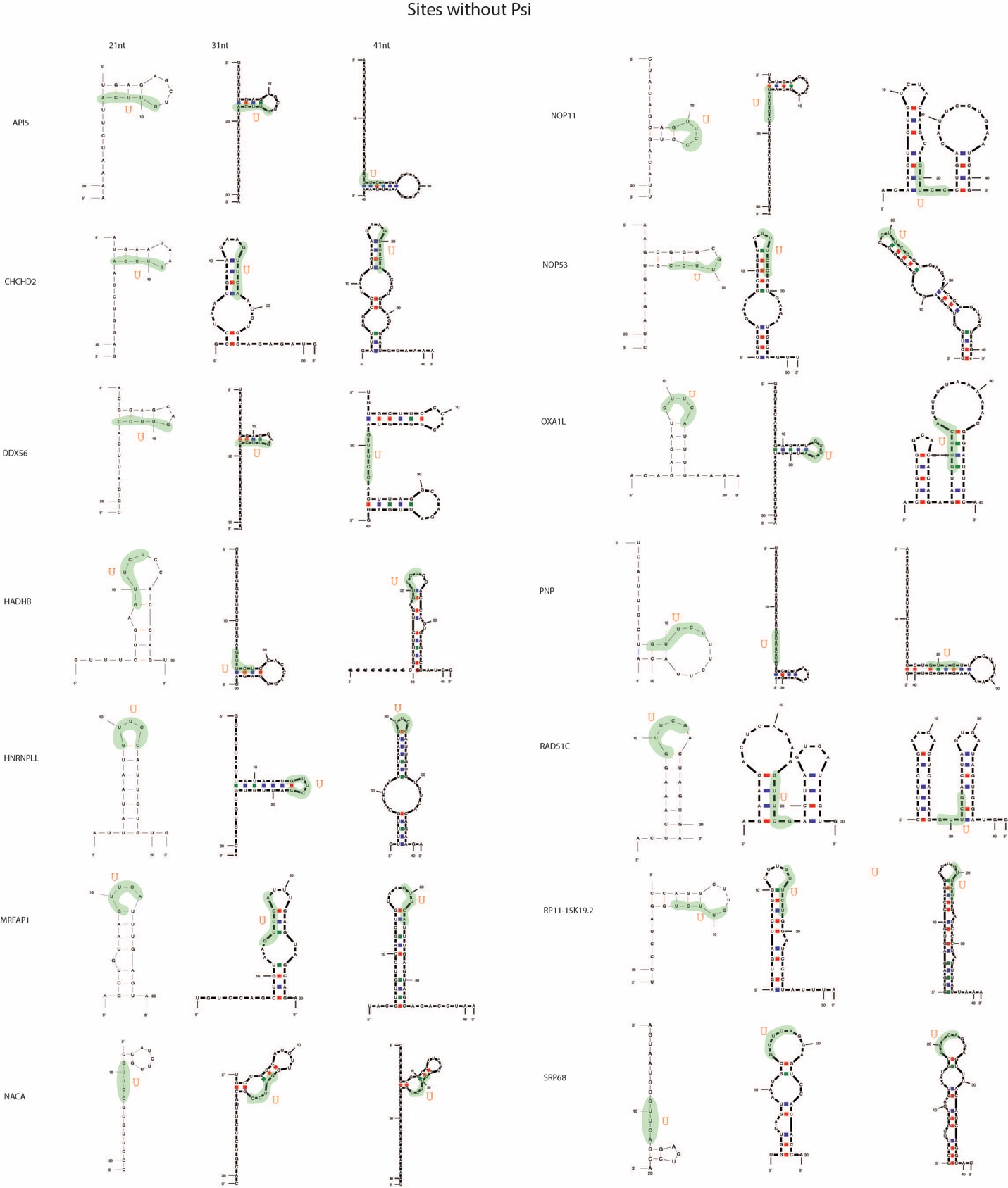

**Supplementary Figure 5.** mFold structures of sites without psi within 21-, 31-, and 41-mer sequence context.

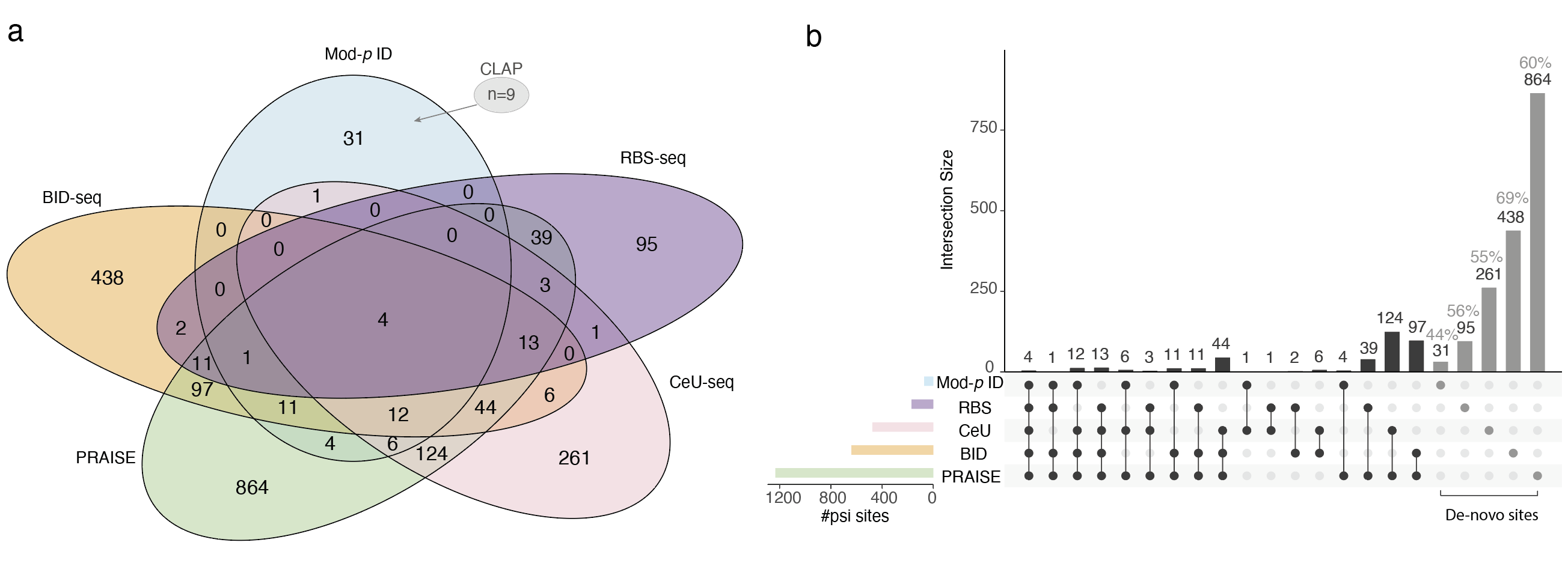

**Supplementary Figure 6.** Comparison of Mod-*p* ID against non-nanopore orthogonal methods for pseudouridine detection.

**a)** A five-way Venn diagram illustrating the overlap of Ψ sites identified by five different methods: Mod-*p* ID (light blue), RBS-seq (purple), CeU-seq (pink), BID-seq (orange), and PRAISE (green). Mod-*p* ID sites used in this analysis are those conserved across all cell types analyzed in this work (see **Supplementary Table S4**). The numbers in each overlapping or non-overlapping region represent the count of Ψ sites identified by the corresponding combination of methods. The grey callout indicates that 9 sites unique to Mod-*p* ID method were previously validated by the CLAP method. b) Intersection of Mod-*p* ID and orthogonal methods. The histogram represents the number of transcripts shared among the indicated cell types. Black dots below each bar indicate which groups contribute to the intersection. Gray bars highlight the de-novo sites identified by each method. The percentages indicate the proportion of each method’s unique sites unconfirmed by other methods.
